# Accounting for salmon body size declines in fishery management can reduce conservation risks

**DOI:** 10.1101/2024.06.06.597779

**Authors:** Jan Ohlberger, Daniel E. Schindler, Benjamin A. Staton

**Affiliations:** School of Aquatic and Fishery Sciences, University of Washington, Seattle, WA 98195; Fish Program, Washington Department of Fish and Wildlife, Olympia, WA 98501; Fishery Science Department, Columbia River Inter-Tribal Fish Commission, 700 NE Multnomah St., Ste. 1200, Portland, OR 97232

**Keywords:** body size, demography, fishing, management strategy evaluation, Pacific salmon, reproductive output

## Abstract

Changes in population demographic structure can have tangible but unknown effects on management effectiveness. Fishery management of Pacific salmon is often informed by estimates of the number of spawners expected to produce maximum sustainable yield (S_MSY_), which implicitly assumes that reproductive output per spawner does not change over time. However, many salmon populations have experienced long-term trends in age, sex, and length-at-age compositions that have resulted in smaller body sizes of mature fish. We present an empirically-based simulation approach for evaluating management implications of declining reproductive output that results from shifting demographics. We simulated populations with or without demographic trends, selective or unselective harvests, and harvest policies based on assessment methods that did or did not account for demographic trends when estimating S_MSY_. A management strategy evaluation showed that expected mean harvests and run sizes were reduced when populations exhibited negative demographic trends. Reduced abundances and increased conservation risks could be partially mitigated by using a stock-recruit analysis based on egg mass instead of spawner abundance, or by using a precautionary management strategy where target escapements were higher than the estimated S_MSY_, especially in fisheries that selectively removed large fish. Accounting for demographic trends in stock-recruit analyses resulted in up to 25% higher run sizes and up to 20% lower conservation risks compared to traditional methods when trends toward smaller, younger, and male-biased runs were present in the population. Conservation of population demographic structure may be critical for sustaining productive fish populations and their benefits to ecosystems and people.

## INTRODUCTION

Pacific salmon (*Oncorhynchus spp*.) are keystone species in freshwater and marine ecosystems and are economically and culturally valuable to people living around the North Pacific Rim. Many populations of Pacific salmon that spawn in freshwater systems of western North America have experienced long-term declines in the average body size of mature fish since at least the 1970s (Bigler et al. 1996; Jeffrey et al. 2017; Losee et al. 2019; Oke et al. 2020). Size declines are particularly pronounced in Chinook salmon (*O. tshawytscha*) (Lewis et al. 2015; Ohlberger et al. 2018, 2019), with differences among populations linked to life-history and ocean distribution (Buckner et al. 2023). Changes in the mean body size of mature fish are a major concern because they may have negative impacts on fishery catches and freshwater ecosystems (Oke et al. 2020). Furthermore, reductions in average reproductive output associated with declines in mean body size (Ohlberger et al. 2020; Malick et al. 2023) have the potential to undermine the sustainability of current fishery management practices, because salmon management is typically focused on the numerical abundance of spawners and does not account for their demographic composition. The conservation and management implications of demographic changes that result from body size declines of salmon remain poorly understood.

The demographic structure of a population integrates the effects of external drivers on individual growth and survival, which can result from environmental variability, species interactions, and exploitation. Trends in population demographic structure therefore reflect many of the short- and long-term impacts of ecological and environmental changes on population productivity. Previous work showed that demographic trends can affect the recruitment and productivity of populations (Shelton et al. 2015; Ohlberger et al. 2022) and that integrating demographic information into statistical stock-recruit analyses can improve the estimation of reference points used in fishery management (Murawski et al. 2001; Wang et al. 2005). The age, sex, and length composition of spawners matters for recruitment because large females have higher reproductive output (Hixon et al. 2014; Barneche et al. 2018).

In Chinook salmon, large females produce more and larger eggs than small individuals (Healey and Heard 1984; Beacham and Murray 1993). Recent declines in mean body size of Chinook salmon are therefore associated with declines in spawner reproductive potential (Ohlberger et al. 2020; Malick et al. 2023). Maternal effects that result in better offspring survival may also contribute to higher recruitment when the spawning population is composed of older females (Green 2008; Ohlberger et al. 2022). Changes in reproductive output due to trends in demography may therefore need to be considered when developing management reference points (Staton et al. 2021) that aim to optimize harvest opportunity and conservation benefits.

Fishery management of Pacific salmon in North America is commonly informed by the principle of maximum sustainable yield (MSY), and management goals are often expressed in terms of escapement goals that are set to achieve the number of spawners on the spawning grounds that maximize long-term yield (Ricker 1954; Beverton and Holt 1957, Clark et al. 2009). Thus, salmon management is focused on the numerical abundance of fish on the spawning grounds and implicitly assumes that each spawner contributes equally to population replenishment and that stock-recruit relationships are stationary such that the mean and variance in recruitment produced by a given number of spawners are constant over time (Walters and Martell 2004). This is evidenced by the rarity with which the demographic composition of spawners is considered when setting management goals. In addition to apparent trends in the demographic composition of the spawning population, stock-recruit relationships may also vary over time due to changes in environmental conditions. Earlier work on Pacific salmon showed that including time-varying parameters in stock-recruitment models can reduce parameter bias and uncertainty (Peterman et al. 2000). However, these analyses were concerned with cyclic changes in productivity due to climate variability, yet long-term trends in population productivity have become increasingly apparent in many populations of Pacific salmon (Peterman and Dorner 2012; Ruggerone and Connors 2015; Malick and Cox 2016).

Here, we use a simulation approach to evaluate the consequences of shifting age, sex, and length (ASL) composition on fishery management performance when informed by models that differ in how they account for demographic trends when estimating biological reference points. The data-driven simulation model quantifies how reproductive output and the number of spawners needed to produce maximum sustainable yield change when declining trends in average body size of mature fish are present in the population. We used the model to quantify bias in the estimation of escapement at MSY when using traditional spawner-recruit analyses based solely on spawner abundance and therefore did not account for trends. We use a management strategy evaluation approach to quantify the effect of different estimation methods on long-term fishery management performance. We parameterized our model using life-history characteristics and observed demographic trends for Chinook salmon. This species has experienced the most pronounced changes in ASL characteristics of mature fish among all species of Pacific salmon (Ohlberger et al. 2018; Losee et al. 2019; Oke et al. 2020). Mean body lengths of adult Chinook salmon in North America have declined by nearly 10% since the 1970s (Ohlberger et al. 2019; Oke et al. 2020) which translates into as much as a 35% decline in reproductive output (i.e., total egg mass) over this time period (Ohlberger et al. 2020). The analysis presented explores the management implications of such size declines and is relevant to the conservation of Pacific salmon throughout their range.

## METHODS

Our approach to evaluating implications of changing demographics for fishery management objectives is illustrated in Figure 1. We considered various scenarios of harvest selectivity, management strategies, and methods for estimating biological reference points that do or do not account for demographic trends. The approach consisted of closed-loop simulations based on an operating model to generate stochastic population data with or without demographic trends, and an estimation model used to fit stock-recruit models to estimate biological reference points.

**Figure 1:**
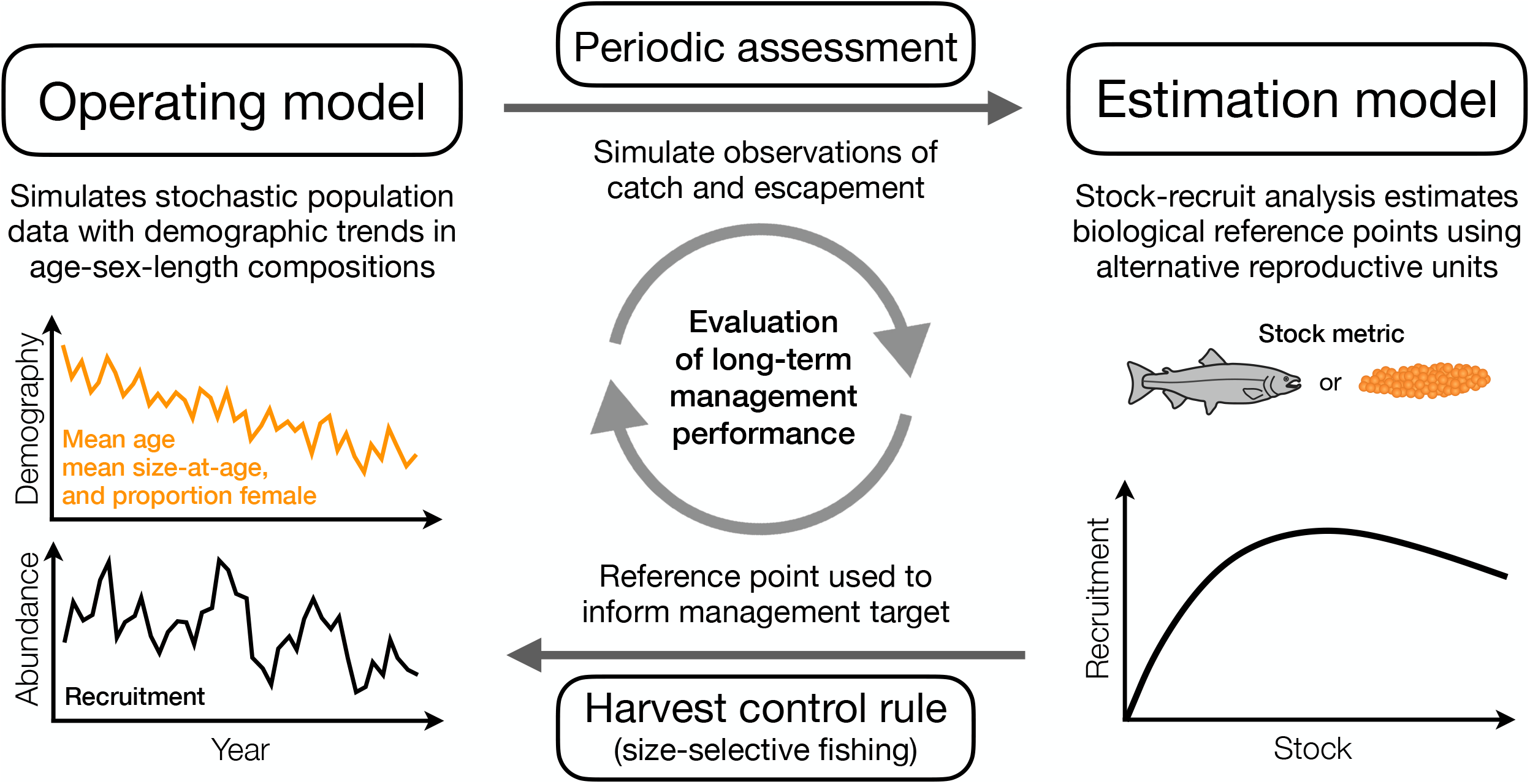
Conceptual illustration of the approach. An operating model was used to simulate stochastic population data with demographic trends in the age-sex-length compositions of recruits that are harvested by a potentially size-selective fishery. Observations of the catch and escapement were then made with error from the simulated recruitment outcomes and passed periodically to an estimation model that used a stock-recruit analysis to estimate biological reference points, in particular the escapement at MSY, using different estimation methods that did or did not account for trends in demographic trends. The estimates inform the harvest control via a management target (e.g., escapement goal), by using the reference point estimate directly or a more liberal or precautionary strategy based on its value. The long-term performance of the fishery management, relative to quantitative utility measures, was evaluated under different scenarios of demographic trends, harvest selectivity, methods for estimating reference points, and management strategies.

Simulated data from the operating model were passed to the estimation model for assessment every ten years, which used either spawner abundance or total egg mass as the metric of spawning stock in a stock-recruit model. The estimated biological reference point from the assessment model (escapement at MSY) was periodically used to set a new management target. Simulations were run over multiple periodic assessments and the performance of the fishery management was evaluated at the end of the simulations after 100 years according to multiple metrics intended to index harvest and conservation objectives.

### Simulating population dynamics (operating model)

We simulated stochastic Chinook salmon population data using a Ricker stock-recruit relationship, where the population was modeled to include trends in the age composition and sex ratio of recruits and the length-at-age of returning mature fish. We simulated observations of fishery catches and escapements, with error, and used those observations in an estimation model to derive estimates of biological reference points, as described in the subsequent section. Fishery catches and escapement were based on an escapement goal, set as the estimated escapement at MSY for most scenarios. Harvests were taken using a specified size selectivity, which was varied as an additional dimension of the analyses given a recent study found sensitivity of reference point estimates to different fishing gears (Staton et al. 2021). The following section provides a detailed description of the stochastic simulations that generated the time series data used in the stock-recruit analyses of the estimation model.

#### Recruitment

We used a Ricker stock-recruit relationship to simulate the expected recruitment given spawner abundance:

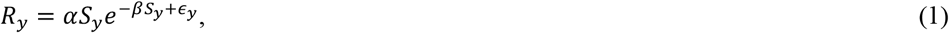

where *R*_*y*_ is the estimated recruitment arising from spawner abundance *S*_*y*_ in brood year *γ, α* represents recruits per spawner as spawner abundance approaches zero, and *β* represents the strength of density dependence. The parameter *β* was drawn from a lognormal distribution:

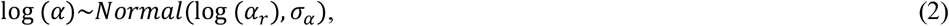

where *α*_*r*_ is the default parameter value and σ_*α*_ is the log-scale standard deviation. The density dependence parameter *β* was also drawn from a lognormal distribution:

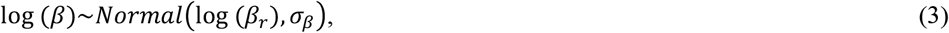

where *β*_*r*_ is the default parameter value and σ_*β*_ is the log-scale standard deviation. The Ricker parameters were adjusted using previously estimated ratios when using egg mass as the stock metric and were applied annually based on the estimated annual reproductive output of spawners (see section ‘Parameter values’ and Staton et al. 2021).

The error term ∈_*y*_ was modeled as autocorrelated residuals (Thorson et al. 2014):

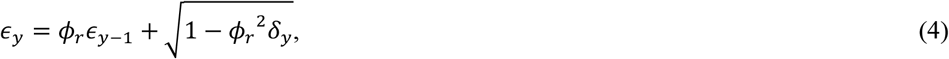

where *ϕ*_*r*_ represents first-order autocorrelation, *ϵ*_*y*_ and *ϵ*_*y*−1_ are recruitment residuals in years *y* and *y* − 1, respectively, and *δ*_*y*_ are normally distributed errors with bias correction for the mean:

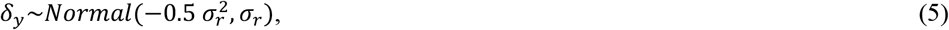

such that σ_*r*_ is the standard deviation of the random component of the recruitment variation. The bias correction ensures that the expected mean value is zero based on a log-normal error variance (Quinn and Deriso 1999).

Recruits (*R*_*y*_) from brood year *y* were apportioned to adult abundances in calendar year *t* by age and sex (*N*_*a,s,t*._) based on the expected sex-specific age-at-maturity proportions of the recruitment (π_*a,s,y*_):

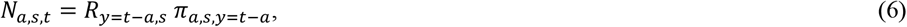

The sex-specific age proportions of recruits from brood year *y* were drawn from a Dirichlet distribution:

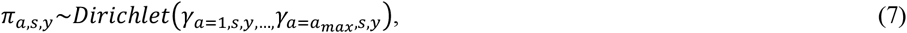

where the shape parameters (*γ*_*a,s,y*_) were based on re-scaled densities derived from a normal distribution around the mean age by sex (*A*_*s,y*_) in brood year *y* with standard deviation σ_*a*_.

Variability in age-at-maturity among brood years was controlled by multiplying the Dirichlet shape parameter vector by *d* (set to 50, smaller numbers would give more variability).

Total abundance in calendar year *t* was taken as the sum of the age- and sex-specific abundances:

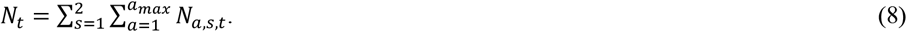

#### Harvest and escapement

Fishery management was guided by an escapement goal. Because escapement goals are not achieved precisely in salmon management due to practical challenges related to run forecasts, run timing uncertainty, mixed-stock considerations, fishing effectiveness, processor capacities, and market dynamics, we assumed that the actual escapement in a given year could differ from the escapement goal target. Spawner escapement in year *t* (S_*t*_) was calculated based on the escapement goal (S_*goal*_) and a log-normal implementation error (*ϵ*_*i*_):

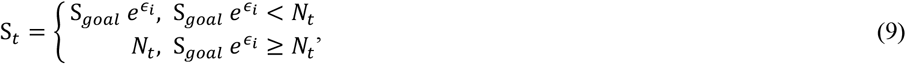

where *ϵ*_i_∼*Normal*(0,*σ*_i_) are normally distributed errors. This formulation ensures that the escapement cannot be larger than the total run in a given year.

Total harvest was calculated as the difference between run size and spawner escapement:

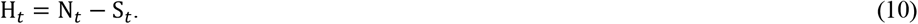

Harvest selectivity was modeled using the Pearson gillnet selectivity model developed for Chinook salmon (Bromaghin 2005), which expresses selectivity as a function of the ratio between fish size and mesh perimeter such that a single set of parameters explains fishery selectivity for all mesh sizes:

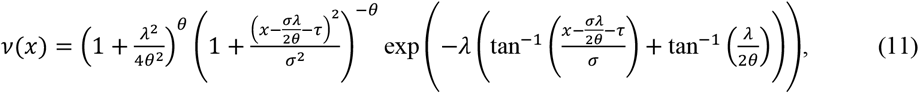

where *x* is the ratio between fish length and mesh perimeter and *λ, θ, σ, τ* are species-specific parameters (Bromaghin 2005; Staton et al. 2021).

Harvest was removed from the run using unequal probability sampling without replacement, based on probabilities of being harvested given age, year, and mesh size (ϖ_*a,t,m*_):

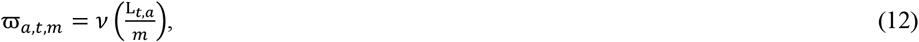

where L_*t,a*_ is mean length of age *a* in year *t*, and *m* indicates the mesh size of the fishing gear. Note that heterogeneity in this vector leads to escapement having a different ASL composition than the run. Using the unequal probability sampling, H_*t*_ was always achieved exactly.

#### Demographic trends

We simulated demographic trends without making assumptions about the causes of these trends. We set default values for the initial mean age, size-at-age, and the proportion of females in the population at the beginning of the simulation based on empirical data for Chinook salmon in western Alaska rivers (Ohlberger et al. 2018, 2020, Staton et al. 2021), and added trends in these demographic characteristics over time. The mean age, mean size-at-age, and proportion female in a given year were drawn from respective distributions to reflect annual variation in demographic characteristics.

Mean age (*A*_*y*_) of returns from a given brood year *y* was drawn from a normal distribution with a mean depending on the initial mean age (*A*_0_) and the change in mean age over time (*δ*_*a*_), and a standard deviation (*σ* _*a*_) describing annual variation around the deterministic trend:

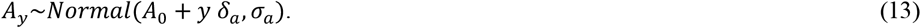

Mean age of female recruits was given by *A*_*f,y*_ = *A*_*y*_ + 0.5 Δ*A*, and mean age of male recruits was given by *A*_*m,y*_ = *A*_*y*_ − 0.5 Δ*A*, where Δ*A* is the mean age difference between the two sexes (females tend to return as older fish compared to males, see parameter values).

The proportion of females (*ψ*_*f,y*_) from a given run year was drawn from a logit-normal distribution with a mean depending on the initial proportion female (*ψ*_*f*,0_) and change in the log-odds of being female over time (*δ*_*f*_), and a standard deviation (*σ*_*f*_) describing annual variation around the deterministic trend:

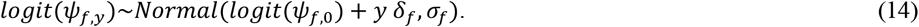

Growth was not modeled explicitly such that size-at-age was provided by run (i.e. calendar) year. Mean length-at-age (*L*_*a,t*_) in a given run year was drawn from a normal distribution with a mean depending on the initial mean length-at-age (*L*_*a*,0_) and a change in mean length-at-age over time (*δ*_*l*_), and a standard deviation (*σ*_*l*_) describing annual variation around the deterministic trend:

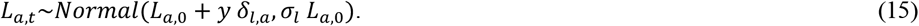

Setting *δ*_*a*_, *δ*_*l,a*_, and *δ*_*f*_ to zero resulted in simulations without any demographic trends (one of the examined scenarios). In addition to simulated changes in demographic characteristics, selective harvest affected the age and size composition of the escapement, though it did not affect the sex composition, because growth and thus size-at-age are not sex-specific in the model.

#### Reproductive output

Reproductive output (*z*) as a function of female length (L) was modeled according to:

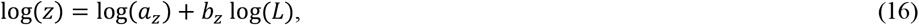

where *z* is reproductive output measured as total egg mass, and *a*_*z*_ and *b*_*z*_ are the intercept and slope of the relationship on the log scale, or the scalar and exponent of the allometric relationship on the arithmetic scale, respectively. The size-scaling of reproduction was assumed to be time-invariant.

Based on the average reproductive output per female spawner by age and year (*z*_*a,y*_), the age-specific proportion of female spawners (*ψ*_*f,a,y*_), and the age-specific spawner abundance (S_*a,y*_) that year, we calculated the total annual reproductive output of all female spawners:

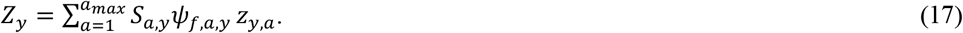

#### Model observations

Simulations made observations of harvest and escapement abundances with error. Observed escapement (*E*_*t*_) was modeled based on true escapement (*S*_*t*_) and lognormal errors:

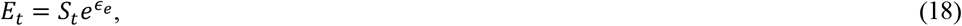

where the error term was bias-corrected just as the white noise portion of recruitment variability in eq. 4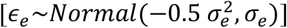. Similarly, observed catch (*C*_*t*_) was modeled based on true harvest (*H*_*t*_) and lognormal errors:

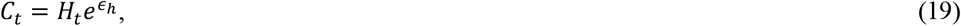

where 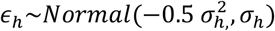.

We assumed that the age compositions of escapement and harvest were observed without error.

#### Parameter values

Parameter values for the stock-recruitment relationship, harvest selectivity, allometric scaling of reproduction, age-sex-length compositions, and demographic trends over time were based on literature values for Chinook salmon. However, different salmon life-histories can easily be implemented by adjusting these values accordingly, and estimates for most parameters are available for other species of Pacific salmon.

##### Stock-recruitment function

To reflect uncertainty in the underlying stock-recruit relationship, Ricker parameters were drawn randomly for each stochastic simulation from lognormal distributions. The parameter *α* is interpreted as the maximum adult recruits produced by one unit of reproductive output and *β* is the inverse of the number of reproductive units expected to produce maximum recruitment. We used mean *α*_*r*_ = 5 and standard deviation *σ* _*α*_= 0.2 for the productivity term and mean *β*_*r*_ = 5*e*^−5^ (i.e. 20,000 spawners produce maximum recruitment) and standard deviation *σ*_*β*_= 0.2 for the density dependence term, based on previous work on Chinook salmon (Fleischman et al. 2013). However, these values only apply to stock-recruit relationships based on spawner abundance, whereas our simulations used total egg mass as the stock metric. We therefore converted parameter values using estimates from a recent empirical study that fitted Ricker models based on spawner abundance (N) or total egg mass (E) and accounted for demographic trends (Staton et al. 2021):

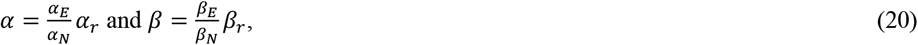

where previously estimated parameter values were *α*_*N*_ = 5.07 or *α*_*E*_ = 0.01036 for productivity and *β*_*N*_ = 8.6*e*^−6^ or *β*_*E*_ = 1.7*e*^−8^ for the density-dependence term (Staton et al. 2021).

Values for the autocorrelation (*ϕ*_*r*_ = 0.4) and standard deviation (*σ*_*r*_ = 0.35) of recruitment residuals were based on previous stock-recruit analyses on Chinook salmon (Staton et al. 2021). However, stronger residual autocorrelation has previously been reported for Chinook salmon (Fleischman et al. 2013), and other species of Pacific salmon may exhibit lower autocorrelation and higher variation in recruitment (Thorson et al. 2014), presumably owing to their less diverse spawner age distributions.

##### Age composition

We modeled Chinook salmon from total age 1 to 9. We assumed that the youngest age group returning to spawn was age 3, after freshwater and ocean residence. The overall mean age at the start of the simulation was set to *A*_0_ = 5.5 years, and the difference between sexes was Δ*A* = 1, such that the average age of males and females without trends was 5 and 6 years, respectively. Shape parameters of the Dirichlet distribution were based on normal probability densities (gamma, eq. 7) around the mean age by sex using a standard deviation of *σ* _*a*_ = 0.6. Consequently, the majority of the return from a given brood year was initially composed of total ages 4, 5, 6, and 7, with males primarily returning at ages 4-6 and females at ages 5-7. The proportional contributions of all other age groups were typically below 1%. This age distribution was representative of high latitude Chinook salmon populations in the 1970/80s (Ohlberger et al. 2018). When included in scenarios, the trend in mean age was set to *δ* _*a*_ = −0.008, i.e., a decline of -0.4 over 50 years, consistent with empirical data on Chinook salmon (Ohlberger et al. 2018; Staton et al. 2021).

##### Size-at-age

Size-at-age distributions at the start of the simulation were based on von Bertalanffy growth parameters describing the marine growth of the fish. We used an average asymptotic length (mm) of L_*inf*_ = 1200, a growth rate coefficient of *k* = 0.325, and an ocean entry length (mm) of *S*_*e*_ = 150, yielding initial mean lengths-at-age (L_*a*,0_) at *t* = 0 of 650, 800, 910, and 990 mm for fish that return after 4, 5, 6, and 7 years, respectively (typical ocean ages 2-5; Ohlberger et al. 2018, 2019). Mean length-at-age in a given run year was drawn from a normal distribution with standard deviation *σ*_*l*_ equal to 0.1 times the initial mean length-at-age (eq. 15). When included in scenarios, the default trends in mean length-at-age (*δ*_*l,a*_) over time differed by age group, with younger ages slightly increasing and older ages declining in length-at-age. Changes in mean lengths for return ages 4, 5, 6, and 7 were set to 0.2, -0.6, -1.2, and -2 mm per year, respectively, which is equivalent to total changes of 10, -30, -60, and -100 mm over a period of 50 years. These values are consistent with an empirical analysis that estimated changes in the average lengths-at-age of Chinook salmon since the 1970s of 3%, -5%, -7%, and -9% for ocean ages 2, 3, 4 and 5, respectively (Ohlberger et al. 2018). Similarly, changes in mean lengths of these age groups over that time period were 5%, -6%, -8%, and -10% for female Chinook salmon in the Kuskokwim River, Alaska (Staton et al. 2021).

##### Mean body size

The resulting overall decline in mean body size due to shifts in age composition and length-at-age, when included in the simulated scenarios, was on average about 65 mm or 8% (a typical mean length in years prior to demographic changes was roughly 800 mm). Depending on the selectivity of the fishing gear, these trends translate into changes in mean length in the escapement (after harvest) of about 60-80 mm or 7.5-10%. The simulated trends match observed declines of 70-80 mm or nearly 10% in the mean body size of Chinook salmon in North America (Ohlberger et al. 2019; Oke et al. 2020).

##### Sex composition

The proportion of females from a given brood year at the beginning of the simulations was set to *π*_*f*,0_ = 0.45 (Ohlberger et al. 2020; Staton et al. 2021). When included in scenarios, we used a trend in the proportion female of *δ*_*f*_ = −0.008 per year on logit scale, which is equivalent to a proportional decline of 0.1 over a period of 50 years, i.e. *π*_*f*_,_50_ = 0.35 (Staton et al. 2021).

##### Reproduction

The length-scaling exponent and scalar (slope and intercept on log-log scale) of total egg mass as a function of female length were set to *a*_*z*_ = 8.71*e*^−12^ and *b*_*z*_ = 4.8, based on previous work on Yukon River Chinook salmon (Ohlberger et al. 2020). Because body mass scales roughly to length cubed (*b*∼3.4), total egg mass per female scales hyperallometrically with body mass in Chinook salmon (Ohlberger et al. 2020). We standardized this allometric relationships to a typical female length of 800 mm with a mean egg mass of 916 g at that size.

##### Fishery parameters

Species-specific values for the parameters of the selectivity function were taken from the literature. We used values of *λ* = −0.547, *θ* = 0.622, 6 = 0.204, *σ* = 1.920 for Chinook salmon (Bromaghin 2005). To simulate unselective fishing, we set *σ* = 10. The implementation error of the fishery was assumed to be *σ*_*i*_ = 0.3.

##### Observations

We assumed that observations of escapement were less certain than observations of harvest. While fishery catches may be misreported, escapement estimates are often based on space and time extrapolations that can introduce considerable error. We used observation errors of *σ* _*e*_ = 0.2 for annual escapements and *σ* _*h*_ = 0.1 for annual harvests.

### Estimating biological reference points (estimation model)

Estimation of biological reference points using the Ricker stock-recruit model was based on the observed harvest and escapement and their age, sex, and length compositions. However, we did not account for observation error in the harvest and escapement in the estimation model because fitting a state-space population model (e.g., Fleischman et al. 2013, Staton et al. 2021) to thousands of stochastic simulations was computationally impractical. We therefore used a regression-based approach to estimate parameters of the stock-recruit relationships, which implicitly assumes that spawner and recruit data are independent and observed with negligible error in addition to the aforementioned assumptions about stationarity and repeatability. While regression-based approaches can suffer from bias in estimates of biological reference points (Walters 1985; Walters and Martell 2004), our preliminary analyses indicated little bias in model parameters and estimated reference points when comparing estimates from stock-recruit models fit to simulated true values to estimates from model fit to observations of the simulated data, within the range of observation error values used in the simulations. We used three different regression-based approaches for estimating biological reference points that varied in how they accounted for demographic time trends: a traditional linearized stock-recruit model (no accounting), a dynamic linear model (implicit accounting), and a yield-per-recruit approach (explicit accounting).

#### Traditional Ricker stock-recruit model

A maximum-likelihood estimation of the linearized Ricker stock-recruitment model based on spawner abundance was used as the *traditional* approach to estimating biological reference points used in salmon management. The traditional Ricker model (TRM) assumed time-invariant parameters such that the demographic characteristics of the population are averaged across all years of the assessed time series. It is important to note, however, that using the time-invariant Ricker model did not imply that the management policy was constant through time because management goals were assessed periodically, which allowed the management to slowly track changes in population productivity even when the assessment model assumed stationarity.

Stock-recruit parameters were estimated by fitting the linearized version of the Ricker model using maximum-likelihood estimation. The reconstructed brood year recruitment (*R*_*y*_) was:

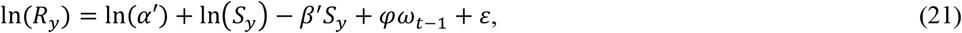

where *S*_*y*_ is spawner abundance in brood year *y, ω*_*y*−1_ is the recruitment residual in year *y* − 1, *φ* is an autocorrelation parameter, and *ε*∼*Normal*(0, *σ*_*r*_) are normally distributed random errors.

The productivity parameter was corrected for the difference between the median and the mean of a lognormal error distribution with a first-order autocorrelation process (Fleischman et al. 2013):

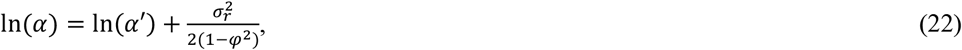

where φ is the autocorrelation and *σ* _*r*_is the standard deviation of the random recruitment error, as described above. Following Hilborn and Walters (1992), the density dependence parameter was corrected as:

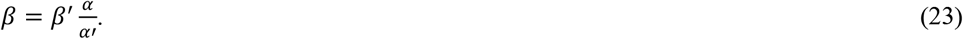

We calculated the escapement that is expected to produce maximum sustainable yield based on the estimated stock-recruit parameters using the explicit solution (Scheuerell 2016):

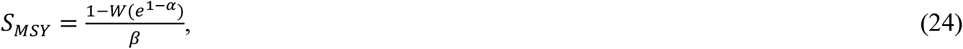

where W is the Lambert function and *α* and *β* are the corrected Ricker parameters.

#### Dynamic Linear Model

A Dynamic Linear Model (DLM) was used to allow for time-varying parameters of the linearized Ricker stock-recruitment model. Here, the *α* and *β* parameters can vary over time to reflect changes in population productivity and density dependence due to trends in the population age, sex, and length compositions. The productivity and density dependence parameters of the linearized Ricker function can be modeled as a function of time as follows:

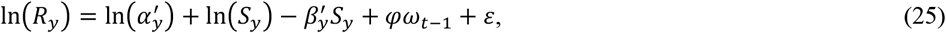

where the time-varying 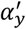 and 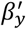 parameters were estimated using maximum likelihood estimation with added Kalman filtering and backward recursive smoothing as implemented in the R package *dlm* (Petris 2009). The estimated productivity parameter was corrected for the difference between the median and the mean of a lognormal error distribution (see eq. 22). Escapement at MSY estimates for the DLM were based on the time-varying stock-recruit parameters as follows:

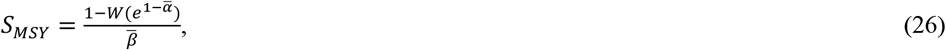

where 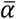 and 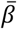 are average parameter values for the most recent five years prior to assessment.

#### Equilibrium calculations for alternative reproductive units

To explicitly account for demographic trends, we used a yield-per-recruit (YPR) approach based on equilibrium calculations to obtain estimates of escapement at MSY (Botsford 1981; Walters and Martell 2004). Here, total reproductive output is used as the stock metric in the Ricker stock-recruit model, instead of spawner abundance. The fishing mortality that maximizes equilibrium harvest is then determined using an optimization algorithm (e.g., Staton et al. 2021) and the resulting equilibrium escapement is taken as the escapement that produces maximum sustainable yield. This yield-per-recruit analysis assumes that the allometric relationship between reproductive output and body size is known and that data on the size composition of spawners are available, such that the expected average reproductive output per spawner can be calculated.

The equilibrium calculations for alternative reproductive units were performed according to equations 12-16 in Staton et al. (2021). An optimization algorithm was used to determine the fishing mortality associated with the highest equilibrium harvest, and to calculate the total equilibrium escapement, which was taken as *S*_*MSY*_. Equilibrium escapement depends on the mesh size used in the fishery because selectivity affects the age-specific probability of escaping harvest. Our reference period for setting new management targets was the last five years of the assessed time series. This means that the equilibrium escapement was based on the average reproductive output-per-recruit over the five years prior to the assessment.

#### Fishery management targets

We used the estimated escapement at MSY to update the escapement target (S_goal_) used in the fishery unless the new estimate was not within 0.5-2 times the previous escapement target. These limits were set to avoid unrealistic targets, especially early in the time series when few years were available to inform the estimation model. We further explored more liberal and precautionary management strategies by using either 0.75 S_MSY_ or 1.5 S_MSY_ as the new management target, respectively. These scenarios were implemented to represent systems in which the managers favor a liberal approach (0.75), i.e. more harvest, or a precautionary approach (1.5), i.e. more escapement. The precautionary approach of using 1.5 S_MSY_ is similar to using an upper target of S_MAX_, the spawner abundance that maximizes recruitment (S_MAX_ equals about 1.5 S_MSY_ across all scenarios in our analysis). S_MAX_-based escapement goals are sometimes used in fishery management and are deemed more appropriate where the fishery is dominated by subsistence and sport fisheries that wish to harvest a fixed number of fish each year and are attempting to minimize the effort needed to harvest (Liller and Savereide 2022). The escapement target (S_goal_) was then updated with the new estimate of S_MSY_, which was subsequently used to dictate fishery catches until the next periodic assessment was performed.

### Assessing fishery management performance

We assessed several different scenarios as part of a management strategy evaluation, each of which included different assumptions about demographic trends, fishery selectivity, estimation method, and the management strategy. Each of these scenarios was evaluated by using 1000 stochastic replicates. We evaluated differences in estimates of the escapement at MSY for alternative estimation methods that accounted for demographic trends compared to the traditional time-invariant method after 50 years of simulations for which trends were parameterized using observed trends in the age-sex-length compositions of North American Chinook salmon.

We evaluated the performance of the fishery management during the following 50 years (after 100 years of simulations), with either (i) no demographic trends, (ii) trends in ASL compositions that stabilized after the historical period for the remaining 50 years, or (iii) ASL trends that continued at the historical rate for the entire simulation period. We used the latter two scenarios to bracket the most likely range of future demographic trends, where stabilized and continued trends in age, sex, and length compositions were considered the most optimistic and pessimistic scenarios, respectively. A scenario where ASL compositions reverse to their historical values was considered unlikely and thus not included. An example of simulated demographic trends and population abundances for a scenario with ASL trends that stabilize after 50 years using a large-mesh selective fishery and a traditional Ricker stock-recruit analysis is shown in Figure S1.

Fishery management performance was evaluated for the last 50 years of the simulations. Metrics used to evaluate management performance were mean annual harvest, mean annual run size, and the probabilities that recruitment and escapement were above certain conservation thresholds.

Mean harvest and mean run size were calculated as the average harvest and run size over the simulated time period. To assess harvest stability, we calculated the inverse of the inter-annual coefficient of variation in harvests, i.e., mean annual harvest divided by the standard deviation in annual harvest. We used abundance thresholds to assess potential conservation risks. First, we calculated the fraction of years that the recruitment (*R*_*y*_) was above 50% of the maximum recruitment of the population (*R*_*max*_):

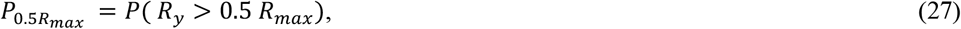

where *R*_*max*_ depends on the stock-recruit parameters as follows (Hilborn and Walters 1992):

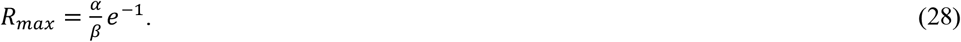

Second, we calculated the fraction of years that the spawner escapement (*S*_*y*_) was above 50% of the equilibrium spawner abundance (*S*_0_):

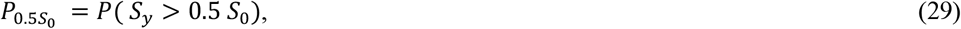

where *S*_0_ depends on the stock-recruit parameters as follows (Hilborn and Walters 1992):

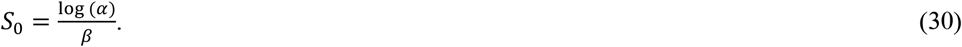

We considered higher mean harvests, higher mean run sizes, and a higher probability of abundance above the conservation thresholds as desirable fishery management objectives.

## RESULTS

Average reproductive output, measured as mean egg mass per spawner, declined over time due to trends in spawner age-sex-length compositions (Figure S2). Declines in mean egg mass per spawner over 50 years were in the 30-45% and 40-55% range (median values) for simulated age-length trends and age-sex-length trends, respectively, depending on the selectivity of the fishery. A large-mesh fishery (8.5-inch gillnets) caused ∼15% stronger declines in body sizes of fish in the escapement than a small-mesh fishery (6-inch gillnets), due to the selective removal of larger individuals (not due to evolutionary changes in response to size-selective fishing, which were not accounted for in the simulations). For reference, the estimated median decline in reproductive output was zero for a population without demographic trends that experienced an unselective fishery. The estimation method had a negligible effect on changes in reproductive output.

Differences in estimates of the escapement that produces MSY (S_MSY_) between the alternative estimation methods depended on the selectivity of the fishery (Figure 2). When demographic trends were present in the population, the Dynamic Linear Model (DLM) resulted in median S_MSY_ estimates that were similar to or slightly lower than the traditional Ricker model (TRM), whereas the yield-per-recruit analysis (YPR) resulted in higher estimates, with the exception of a small-mesh selectivity where estimation method had little effect on S_MSY_. For a large-mesh selective fishery, the YPR method resulted in S_MSY_ estimates that were about 24% higher for age-length trends and about 30% higher for age-sex-length trends (median values) compared to the TRM. For an unselective fishery, the median % difference was about 11% and 18% for age-length and age-sex-length trends, respectively. Under a small-mesh selective fishery, the YPR method resulted in S_MSY_ estimates that were lower than estimates of the TRM, especially when no demographic trends were present in the population.

**Figure 2:**
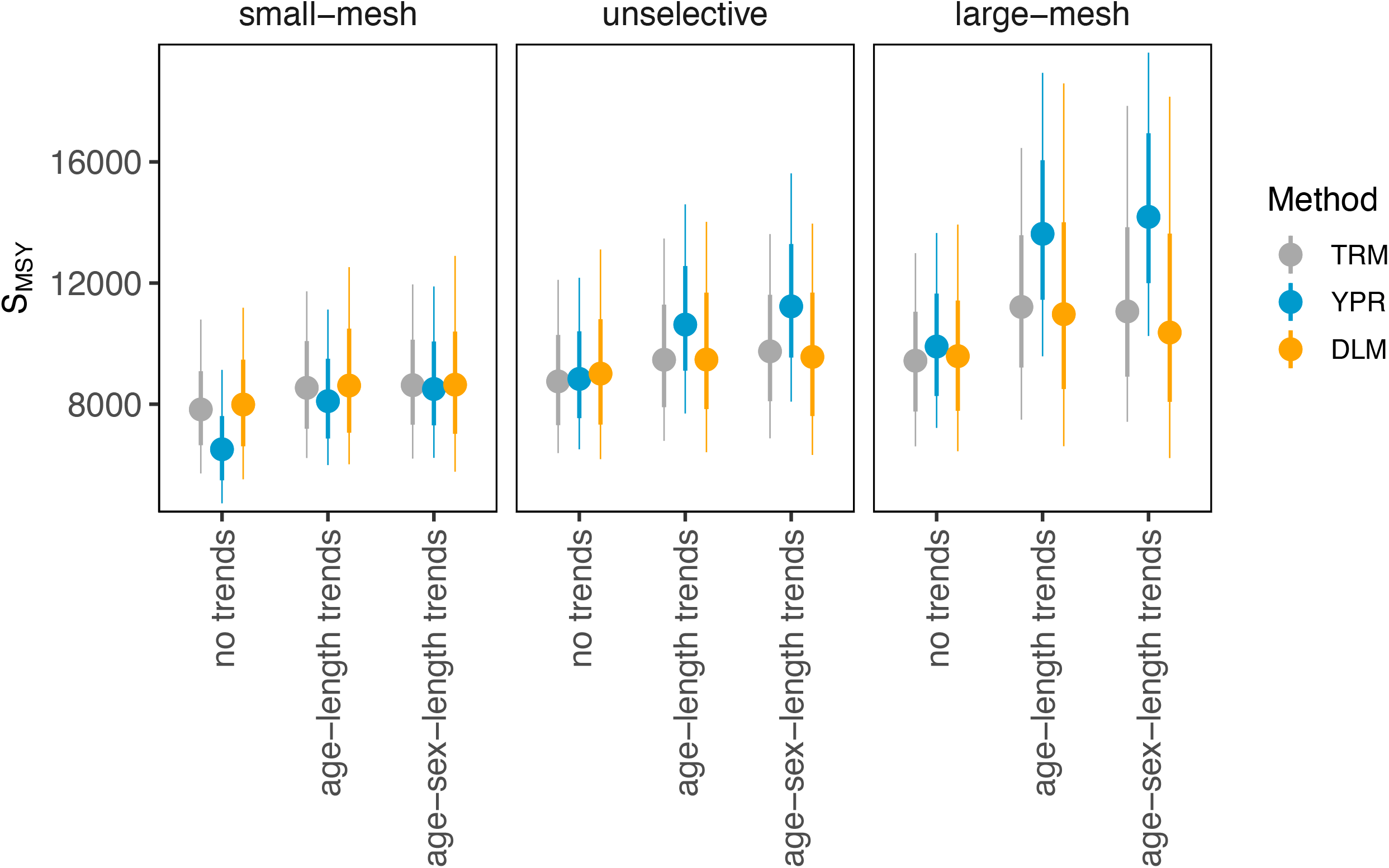
S_MSY_ estimates by estimation method. Shown are median values (circles) with 50% quantiles (thick bars) and 80% quantiles (thin bars) for the traditional Ricker model (TRM, gray), the yield-per-recruit analysis (YPR, blue) and the Dynamic Linear Model (DLM, yellow). S_MSY_ estimates were evaluated at the end of the historical period after 50 years of demographic trends.

The performance of fishery management depended on the estimation method, selectivity of the fishery, and the presence of demographic trends (Figure 3). Mean harvest, mean run size, and the probability that the recruitment was above 50% of the maximum recruitment all declined when demographic trends were present. This overall performance decline is attributable to the reduced reproductive output and associated decline in population productivity. In addition, the probability that the escapement was above 50% of the equilibrium spawner abundance (*S*_0_) declined unless demographic trends were explicitly accounted for using the yield-per-recruit approach.

**Figure 3:**
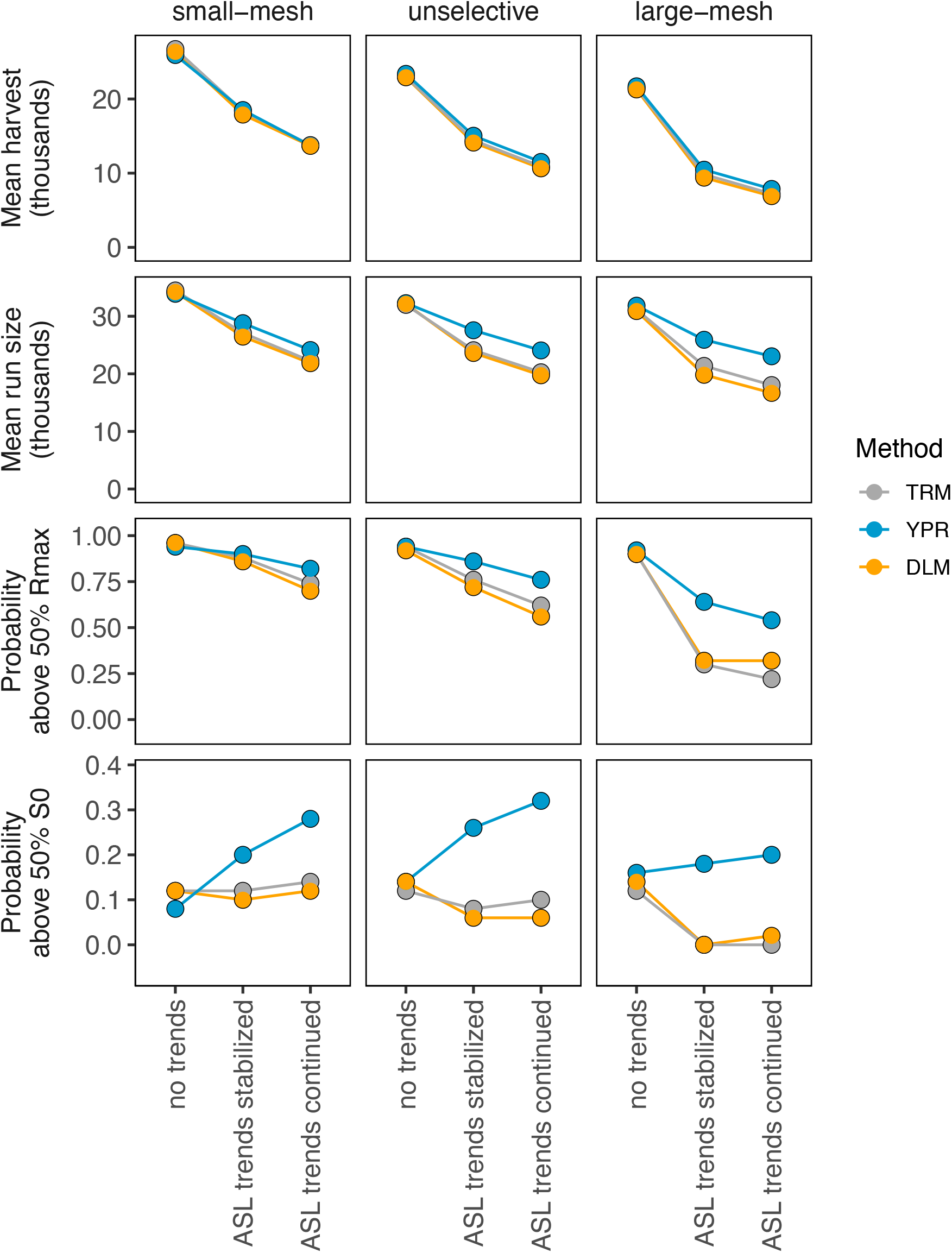
Evaluation of fishery management performance under different scenarios. Shown are median values of four performance metrics (rows) under three different selectivity regimes (columns) for each estimation method (colors) as a function of the demographic trends. Evaluated performance metrics were mean harvest, mean run size, probability that recruitment was above 50% of the maximum recruitment (R_max_), and the probability that the spawner escapement was above 50% of the equilibrium spawner abundance (S_0_). Simulated selectivity regimes were a small-mesh (6-inch), unselective, or large-mesh (8.5-inch) fishery. The three estimation methods were the traditional Ricker model (TRM, gray), the yield-per-recruit analysis (YPR, blue), and the Dynamic Linear Model (DLM, yellow). Age-sex-length (ASL) trends were either not included, stabilized after 50 years, or were assumed to continue into the future (x-axis). Simulations were run for 100 years, and fishery performance was evaluated for the last 50 years.

Compared to the TRM, the YPR method increased the probability that escapement was above 50% of *S*_0_ by up to 0.2, which implies a reduction in the risk of falling below that abundance threshold of 20%. All performance metrics further tended to be lower for a large-mesh selectivity because a fishery that selectively removes large individuals further exacerbates the decline in mean reproductive output per spawner. Moreover, the method used for estimating S_MSY_ had a relatively small effect on harvest such that the three estimation methods resulted in similar mean annual harvests.

The YPR approach clearly outperformed the other estimation methods in terms of mean run sizes and the probabilities that recruitment and escapement were above the conservation thresholds. Specifically, when assuming continued age-sex-length trends, the median percent difference in mean run abundances between the YPR and TRM was about 8%, 17%, and 25% for a small-mesh, unselective, and large-mesh fishery, respectively (Figure 4). These differences between the alternative approaches and the traditional model became more pronounced over time (Figure 5). Furthermore, differences in mean run size were mostly attributable to differences in the mean escapement, rather than differences in mean harvest (Figure 6).

**Figure 4:**
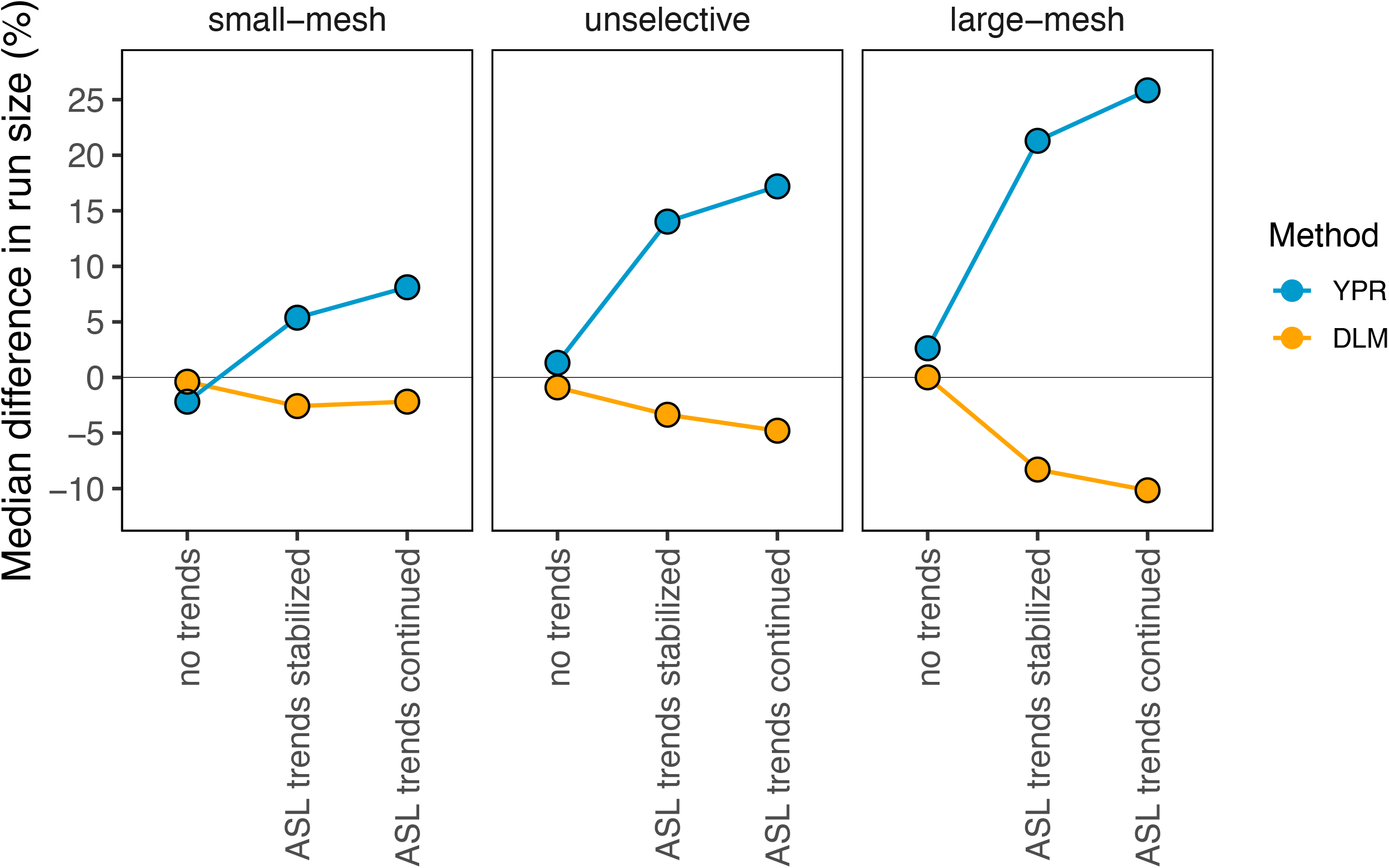
Median % difference in average total run size from the time-invariant model. Shown are differences for the two alternative estimation methods, the yield-per-recruit analysis (YPR, blue) and the Dynamic Linear Model (DLM, yellow), compared to the time-invariant Ricker model for three different fishery selectivities (columns) given the simulated demographic trends (x-axis). This figure is based on numbers presented in the second row of Figure 3.

**Figure 5:**
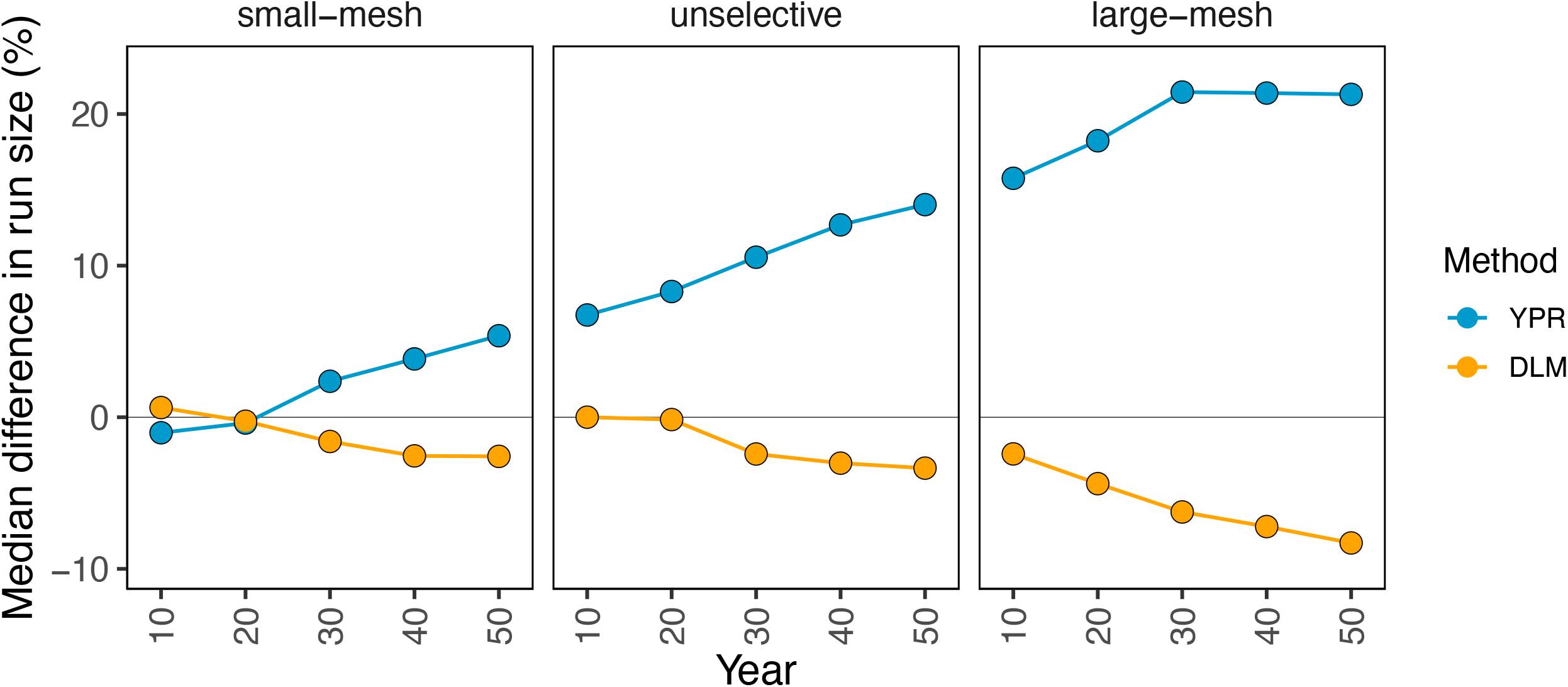
Median % difference in run size from the time-invariant model across years. Shown is the median difference for the yield-per-recruit analysis (YPR, blue) and the Dynamic Linear Model (DLM, yellow) compared to the time-invariant Ricker model in each decade starting after the historical period (e.g., year 10 is the difference for future years 1-10) assuming continued ASL trends.

**Figure 6:**
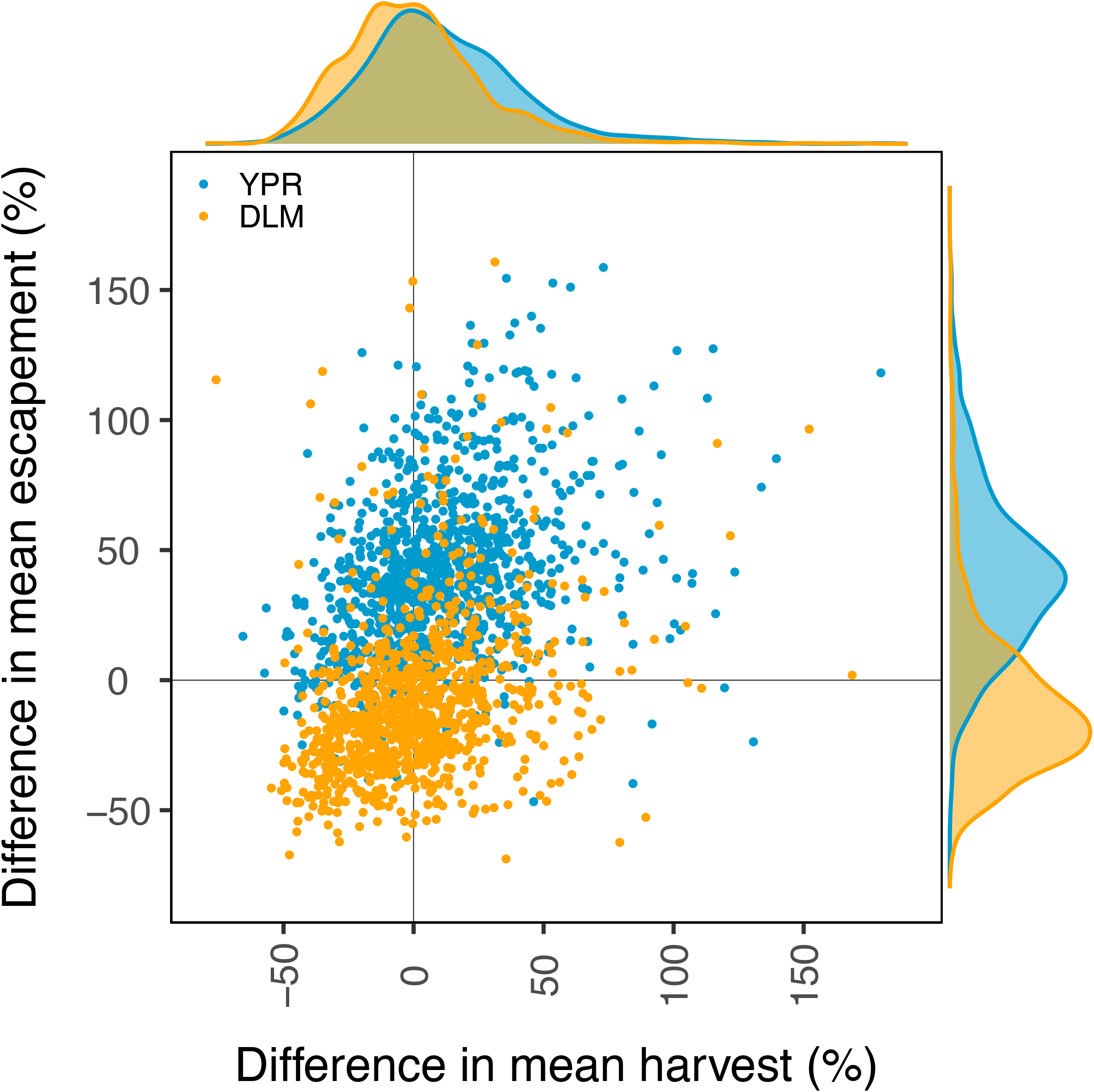
Difference in mean harvest and mean escapement from time-invariant model. Shown is the % difference in harvest and escapement in each stochastic run (circles) for the yield-per-recruit analysis (YPR, blue) and the Dynamic Linear Model (DLM, yellow) compared to the time-invariant Ricker model for a large-mesh fishery assuming continued ASL trends. Marginal plots along the secondary axes show the respective density distributions.

Differences between estimation methods were less pronounced when using a more precautionary management strategy where the escapement target was set to 1.5 times the estimated escapement at MSY (Figure 7). By contrast, when using a more liberal management strategy where the escapement target was set to 0.75 times the estimated escapement at MSY, differences between estimation methods were more pronounced. When assuming continued age-sex-length trends and an unselective fishery, for example, the median percent difference in mean run sizes between the yield-per-recruit and time-invariant approach was about 17% and 25% for the precautionary and liberal management strategies, respectively. For comparison, the ratio of the estimated S_MSY_ for the yield-per-recruit over the time-invariant approach reached a factor of 1.5 or higher only in case of a large-mesh selective fishery. Median values were within the 0.8-1.25 range across all scenarios, which suggests that a slightly lower S_MSY_ multiplier may be sufficiently precautionary (Figure S3).

**Figure 7:**
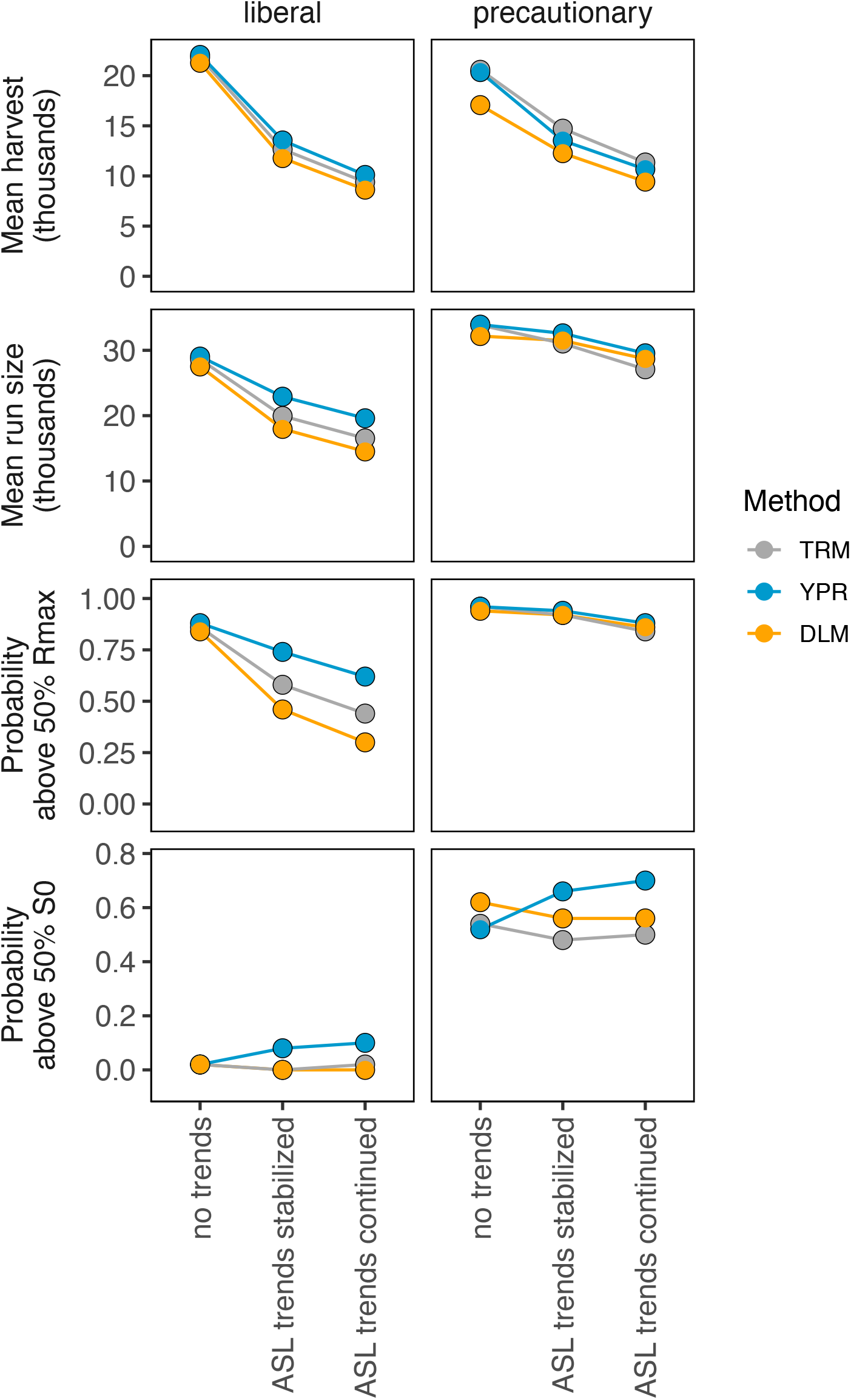
Performance metrics for liberal and precautionary management strategies. Shown are median values of four performance metrics (rows) for each estimation method (colors) using management targets of either 0.75 S_MSY_, (left, liberal) or 1.5 S_MSY_, (right, precautionary), assuming unselective fishing. Details on the performance metrics, the estimation methods, and time period are provided in the caption to Figure 3.

## DISCUSSION

We investigated fishery management implications of demographic trends in age, sex, and length compositions of salmon populations using a broadly applicable data-based simulation approach. We found that compared to a traditional Ricker stock-recruitment model (TRM) based on spawner abundance, and compared to a dynamic linear model (DLM) with time-varying Ricker parameters, an alternative yield-per-recruit (YPR) approach that explicitly accounts for changes in spawner reproductive output results in higher estimates of S_MSY_ in the presence of negative demographic trends. The estimation method affected the fishery management performance when escapement targets were directly based on S_MSY_ estimates. Mean run sizes were higher and conservation risks were lower when using the explicit yield-per-recruit method, while fishery catches were largely unaffected by the choice of model. This pattern was apparent when fishing was unselective and was most pronounced for a large-mesh fishery, in line with previous results (Staton et al. 2021). We found that lower mean run sizes, and higher conservation risks due to demographic trends could also be mitigated notably by using a precautionary management strategy where the escapement target was 1.5 times higher than the estimated S_MSY_. Taken together, our results show that accounting for declines in spawner reproductive output in stock-recruit analyses, or using a precautionary management strategy, can partially mitigate declines in population abundance and reduce conservation risks, especially in fisheries that selectively remove large fish.

### Simulated and observed demographic trends

The demographic trends in age and length compositions used in our simulations are based on observations for Chinook salmon populations along the west coast of North America (Ohlberger et al. 2018, 2019), and trends in sex ratio are based on one of the historically largest populations in North America (Staton et al. 2021). The associated declines in average spawner reproductive output in our simulations (Figure S2) are in line with available empirical estimates for Chinook salmon populations. Reproductive output declined by about 28% for the Yukon River population based on age-length trends (Ohlberger et al. 2020), and by about 49% for the Kuskokwim River population based on trends in age, sex, and length compositions (Staton et al. 2021). While demographic trends over the past few decades have been most pronounced in Chinook salmon, weaker trends toward younger and smaller fish at maturation have also been reported for other species of Pacific salmon, including chum (*O. keta*), coho (*O. kisutch*), and sockeye (*O. nerka*) and pink (*O. gorbuscha*) salmon (Jeffrey et al. 2017; Losee et al. 2019; Oke et al. 2020; Ohlberger et al. 2023).

Shifts in population demographic structures have been linked to changes in environmental and ecological conditions in the ocean such as climate change, intensified predation, and competition with other salmon (Ohlberger et al. 2019; Oke et al. 2020; Manishin et al. 2021). In this study, we did not make specific assumptions about the causes of demographic trends. This also implies that potential evolutionary changes in traits related to growth and maturation in response to size-selective fishing were not accounted for in the simulations. Several studies have concluded that fisheries were not a major driver of recent size declines in Pacific salmon (Kendall at al. 2014; Lewis et al. 2015; Jeffrey et al. 2017; Ohlberger et al. 2019; Oke et al. 2020). Finally, we note that the model does not account for intraspecific density effects on individual growth and hence length-at-age as has been shown for numerically abundant sockeye salmon (Ohlberger et al. 2022), although effects of intraspecific competition on survival are accounted for via the stock-recruit relationship.

### Time-varying productivity models

Previous work that explored time-varying productivity (*α*) in Ricker stock-recruit models suggested that estimates of S_*M*S*Y*_ decrease when population productivity declines, and vice versa (Peterman et al. 2000, 2003). Management targets based on such estimates may thus not be consistent with conservation goals as these models suggest lowering escapement targets when population productivity declines (Holt and Michielsens 2020). However, estimates of *S*_*MSY*_ may not necessarily decline with productivity when the density dependence (*β*) in the stock-recruit relationship also changes over time. For example, assuming a constant maximum recruitment (*R*_*max*_) instead would imply an increase in *S*_*MSY*_ with declining population productivity. To avoid the implicit assumption of stationarity in density dependence, the DLM used in this study allowed for both Ricker parameters to vary over time. This approach nevertheless tended to estimate a lower *S*_*MSY*_ compared to the time-invariant model (consistent with Peterman et al. 2000, 2003). Accordingly, our results suggest that conservation risks may in fact increase when adjusting management targets based on a time-varying Ricker model which may misinterpret declining reproductive potential as declining productivity.

By contrast, explicitly accounting for changes in average reproductive output using the yield-per-recruit method resulted in higher estimates of escapement at MSY and thus leads to management that is better aligned with conservation goals. The yield-per-recruit method resulted in the highest escapement at MSY estimates and the best management performance in terms of mean run sizes and the probability of that run abundance was above the conservation threshold; notably, harvest was not negatively affected with the use of the yield-per-recruit approach.

### Assumptions and limitations of the approach

The yield-per-recruit method has limitations and makes important assumptions about the biology of the population and the data available to the stock assessment. First, this approach assumes that total reproductive output is limited by the number of females but not males. Female spawners are thought to limit population recruitment because eggs are more costly to produce than sperm, spawner sex ratios are often male-biased due to their shorter ocean residence and higher marine survival rates (Olsen et al. 2006; Holtby and Healey 1990), and females defend their nest sites without assistance from males (McPhee and Quinn 1998). Second, this approach assumes that the length composition of the spawners is known. Such information is available for some but not all salmon populations. Third, the yield-per-recruit approach also assumes that the relationship between female size and reproductive output is known. While this assumption may seem limiting due to lack of population-specific data in some systems and potential differences in the allometry of reproductive output among populations (Healey and Heard 1984), such variation is small compared to the implicit assumption of a traditional abundance-based stock-recruitment model that reproductive output does not depend on body size. The length scaling exponent used in this study was 4.8, which is roughly equivalent to a mass scaling exponent of 1.3 in Chinook salmon (Ohlberger et al. 2020). This value is typical for marine fishes and reflects the commonly found hyper-allometric relationship between reproductive output and body size (Barneche et al. 2018). A recent study on Chinook salmon in Alaska found that varying the allometric exponent in a yield-per-recruit analysis, such as the one used in this study, had a relatively small effect on estimates of biological reference points within the range of observed size-scaling relationships (Staton et al. 2021, Supplement A therein). Egg mass was used in this study as a proxy of female reproductive output because it tightly correlates with egg energy content (Ohlberger et al. 2020). Total egg mass may be a better proxy for female reproductive output than egg number, because a trade-off between egg number and egg mass likely evolved to maximize reproductive success in the environment experienced by a population (Heath et al. 2003). Large females produce not only more but also larger eggs compared to small females (Healey and Heard 1984; Ohlberger et al. 2020). Despite the limitations of the yield-per-recruit method, it was the only approach among those tested that allowed explicit accounting of demographic trends and translating them into biological reference points used in fishery management. It should also be noted that we assumed non-retention mortality from small mesh gillnets was negligible for larger fish that interact with the gear but are not captured, yet studies have indicated that gillnet injuries may cause substantial non-retention mortality in salmon (e.g., Baker and Schindler 2009). Finally, we did not assume compensatory increases in survival and thus spawner abundance, which could partially offset declines in reproductive output per spawner, because observational data did not indicate that marine survvial of Chinook salmon has increased over the examined period.

### Implementation uncertainty of the fishery

Differences in the performance of fishery management among estimation methods used in the assessment model may be partially masked by uncertainties in the harvesting process (e.g., implementation error) that cause the achieved harvest and escapement to deviate substantially from the management target (Eggers and Rogers 1987; Dorner et al. 2009; Collie et al. 2012). Such outcome uncertainty may stem from multiple interacting processes, including errors in pre- or in-season forecasts of run abundance, changes in catchability, non-compliance of harvesters to regulations, and slow adjustments in the capacity of the fishing fleet or processors to changes in population abundance (Dorner et al. 2009; Holt and Peterman 2006; Holt and Folkes 2015).

Previous simulation studies showed that this uncertainty can have large effects on the expected management performance when population productivity varies over time (Dorner et al. 2009). Our simulations assume random log-normal error around the escapement target to account for outcome uncertainty. However, we did not include an implementation bias (Peterman et al. 2000), and the uncertainty associated with the harvesting process likely varies among fisheries, species, and regulatory frameworks. Stock-specific applications of assessment models that incorporate information on changes in spawner reproductive potential should thus consider historical patterns in outcome uncertainty of the fishery or investigate how reducing that uncertainty could affect the expected management performance. Briefly, we found that reducing the implementation error (6_4_, from 0.3 to 0.15) did not affect the relative performance of the estimation methods (Figure S4) and hence would not alter any of our conclusions.

## CONCLUSIONS

Salmon management is commonly governed by the principle of sustainable yield, and quantities such as S_MSY_ assume that all spawners contribute equally to recruitment. Under this approach, long-term declines in average reproductive output per spawner result in continuously declining escapement goals when goals are assessed periodically. Using methods that estimate time-varying productivity in spawner abundance-based assessments further accelerates the process and thus increases conservation risks. In contrast, explicit accounting for reduced spawner reproductive output in salmon stock assessments by using alternative reproductive units, or implicit accounting by implementing a precautionary management strategy, can partially mitigate the increased conservation risks due to trends toward younger, smaller, and male-biased returns. However, because such strategies cannot fully compensate for expected declines in run abundances and associated risks, preserving population demographic structure may be an important conservation strategy for sustaining productive salmon populations and their benefits to ecosystems and people.

To conserve population demographic structure, we need a better understanding of the causes of demographic change, which requires monitoring populations for shifts in age-sex-length compositions and reproductive output. The yield-per-recruit approach assumes that data on size compositions are available to calculate the expected average reproductive output per spawner. In the absence of such data needed to explicitly account for changes in demographic parameters, managing harvest in a more precautionary manner, as exemplified by using 1.5 times S_*M*S*Y*_ as the management target, can ameliorate much of the reductions in management performance. The precautionary approach showed little effect on long-term average harvests but improved long-term conservation outcomes. While we did not consider effects of precautionary management on other conservation goals such as maintaining population diversity in large rivers (Connors et al. 2020), such approaches would also work to improve those outcomes.

## ACKNOWLEDGEMENTS

Funding for this work was provided by the North Pacific Research Board (NPRB). We thank Thomas Buehrens for helpful discussions and feedback on previous versions of this work.

## DATA AVAILABILITY STATEMENT

Code for this analysis is available at https://github.com/janohlberger/SizeShiftsImplications.

---

**Table 1.** Parameters values for simulating stochastic population data with demographic trends.

| Symbol | Description | Value | Source |
| --- | --- | --- | --- |
| ***Ricker stock-recruitment relationship*** |  |  |  |
| $\alpha_r$ | Mean Ricker productivity parameter | 5 | 1, 2 |
| $\beta_r$ | Mean Ricker density dependence parameter | $5e^{-5}$ | 1, 2 |
| $\sigma_r$ | Standard deviation in recruitment residuals | 0.35 | 1, 2 |
| $\phi_r$ | Autocorrelation in recruitment residuals | 0.4 | 2 |
| $\sigma_\alpha$ | Standard deviation in $\alpha$ (log scale) | 0.2 | 2 |
| $\sigma_\beta$ | Standard deviation in $\beta$ (log scale) | 0.2 | 2 |
| ***Demographic characteristics and trends*** |  |  |  |
| $A_0$ | Mean age at beginning of simulation | 5.5 | 3 |
| $\sigma_a$ | Standard deviation in mean age (log scale) | 0.6 | 3 |
| $\delta_a$ | Annual trend in mean age | -0.008 | 3 |
| $\Delta A$ | Difference in mean age between sexes | 1.0 | 3 |
| $\pi_{f,0}$ | Initial proportion female | 0.45 | 1, 5 |
| $\delta_f$ | Annual trend in proportion female (logit scale) | -0.008 | 1, 5 |
| $L_{inf}$ | Average asymptotic length (mm) | 1200 | 4 |
| $k$ | Growth rate coefficient | 0.325 | 4 |
| $L_s$ | Length at ocean entry (mm) | 150 | 4 |
| $\sigma_l$ | Standard deviation multiplier mean size-at-age | 0.01 | 3 |
| $\delta_{l,a}$ | Trends in mean size-at-age ($\text{mm} \cdot \text{yr}^{-1}$) | 0.2, 0.6, -0.6, -1.2, -1.8 | 3 |
| $b_z$ | Slope of allometric relationship (egg mass) | 4.8 | 5 |
| $a_z$ | Intercept of allometric relationship (egg mass) | $8.71e^{-12}$ | 5 |
| ***Fishery*** |  |  |  |
| $\lambda$ | Harvest selectivity parameter | 0.547 | 6 |
| $\theta$ | Harvest selectivity parameter | 0.622 | 6 |
| $\tau$ | Harvest selectivity parameter | 1.920 | 6 |
| $\sigma$ | Harvest selectivity parameter | 0.204 or 10 | 6 |
| $m$ | Mesh size in inches | 6.0 or 8.5 | 6 |
| $\sigma_i$ | Fishery implementation error | 0.3 | - |
| ***Observations*** |  |  |  |
| $\sigma_e$ | Observation error on escapement | 0.2 | - |
| $\sigma_h$ | Observation error on harvest | 0.1 | - |

**Sources:** 1) Fleischman et al. 2013; 2) Staton et al. 2021; 3) Ohlberger et al. 2018; 4) Ohlberger et al. 2019; 5) Ohlberger et al. 2020; 6) Bromaghin 2005

## SUPPLEMENRATY INFORMATION

**Figure S1:**
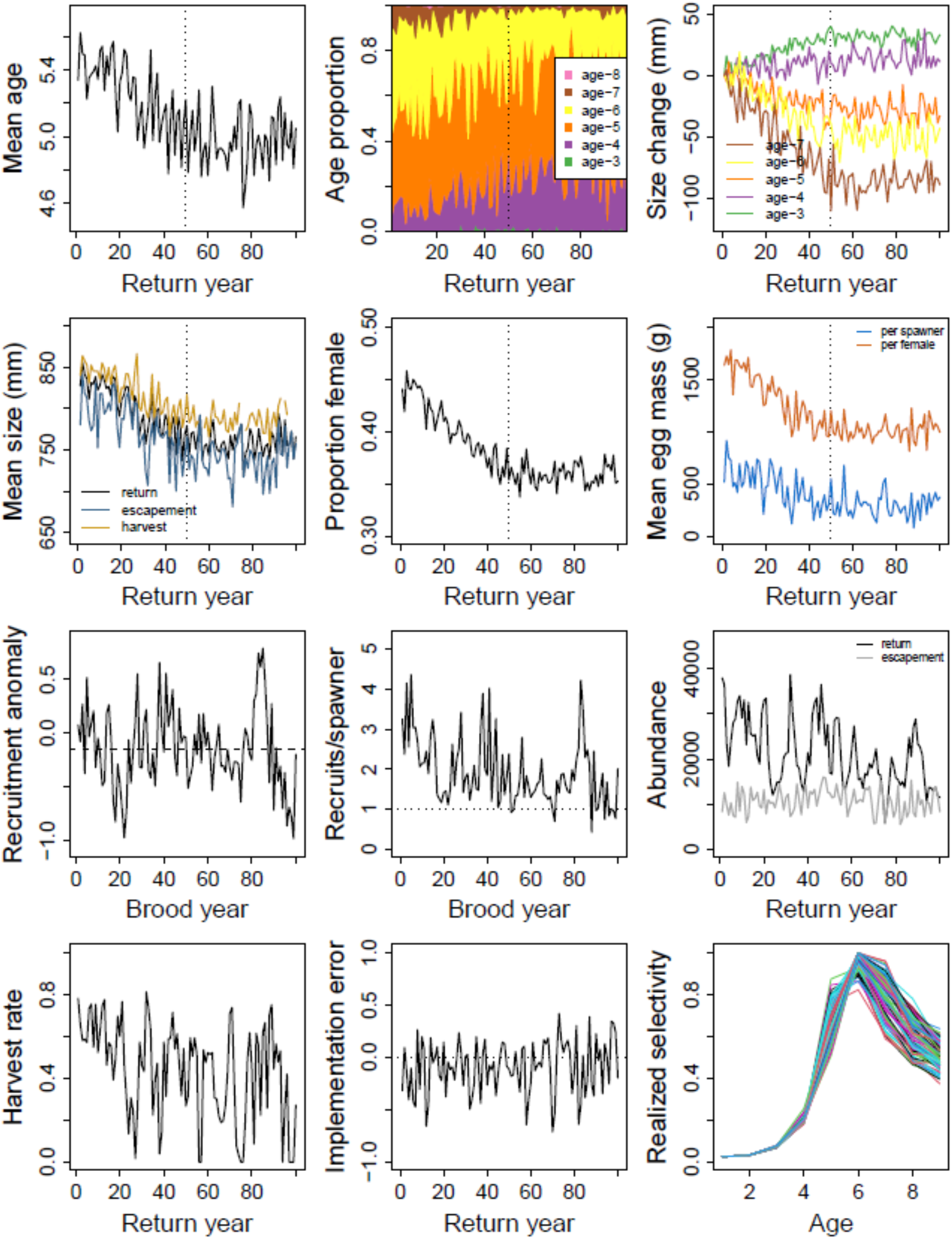
An example of simulated demographic trends and population abundances. Shown is the scenario with age-sex-length trends that stabilize after 50 years, a large-mesh selective fishery, and harvest management based on an S_MSY_ escapement goal that was estimated using a traditional Ricker stock-recruit analysis.

**Figure S2:**
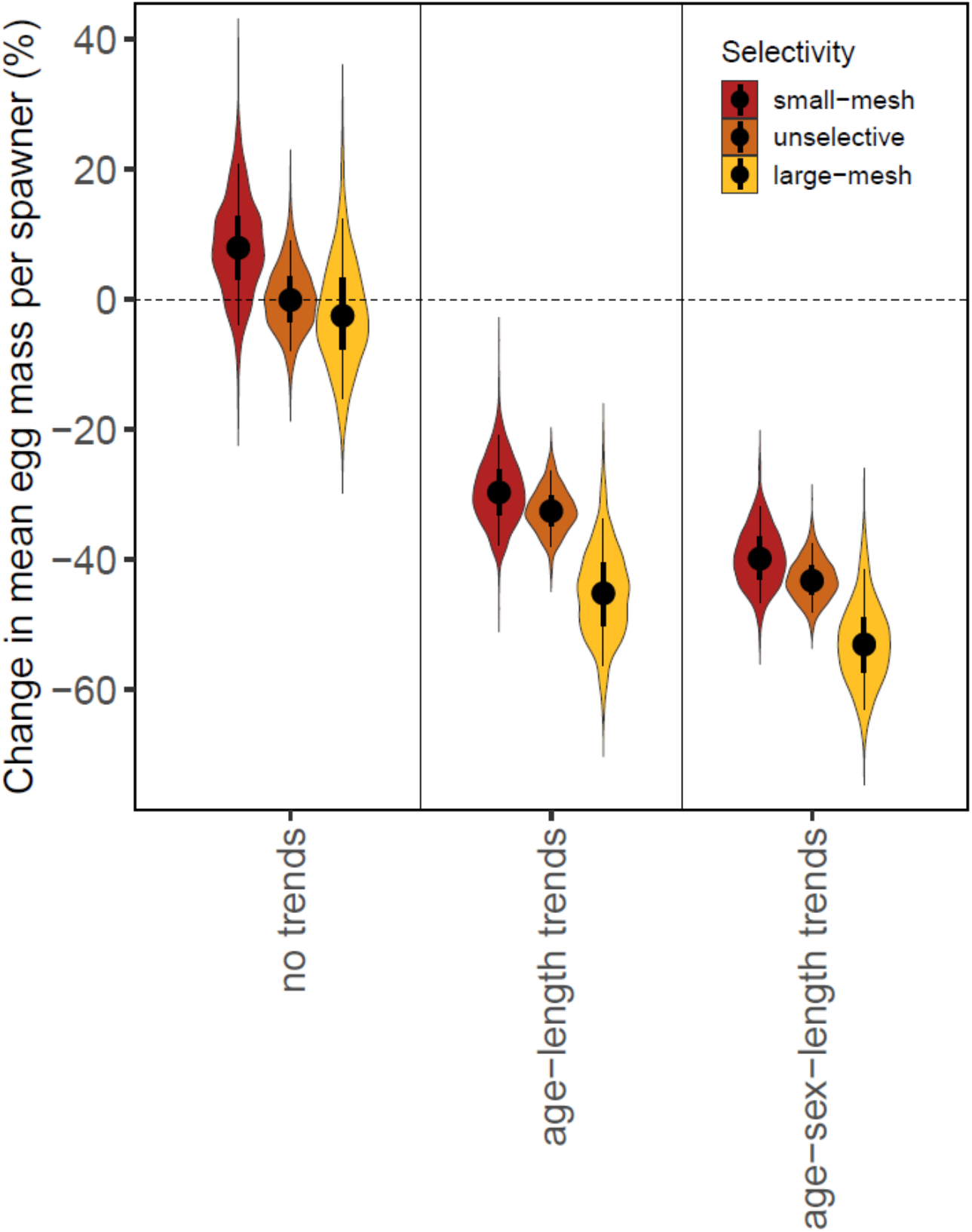
Simulated changes in mean egg mass per spawner over a period of 50 years. Shown are changes over time from the early to late period (first and last ten years of time series) for three trend scenarios: no demographic trends (left), trends in age and length compositions (center), or trends in age, sex, and length compositions (right). Different fishery selectivity types are shown as colors representing a small-mesh fishery (6-inch gillnets, red), an unselective fishery (orange), and a large-mesh fishery (8.5-inch gillnets, yellow).

**Figure S3:**
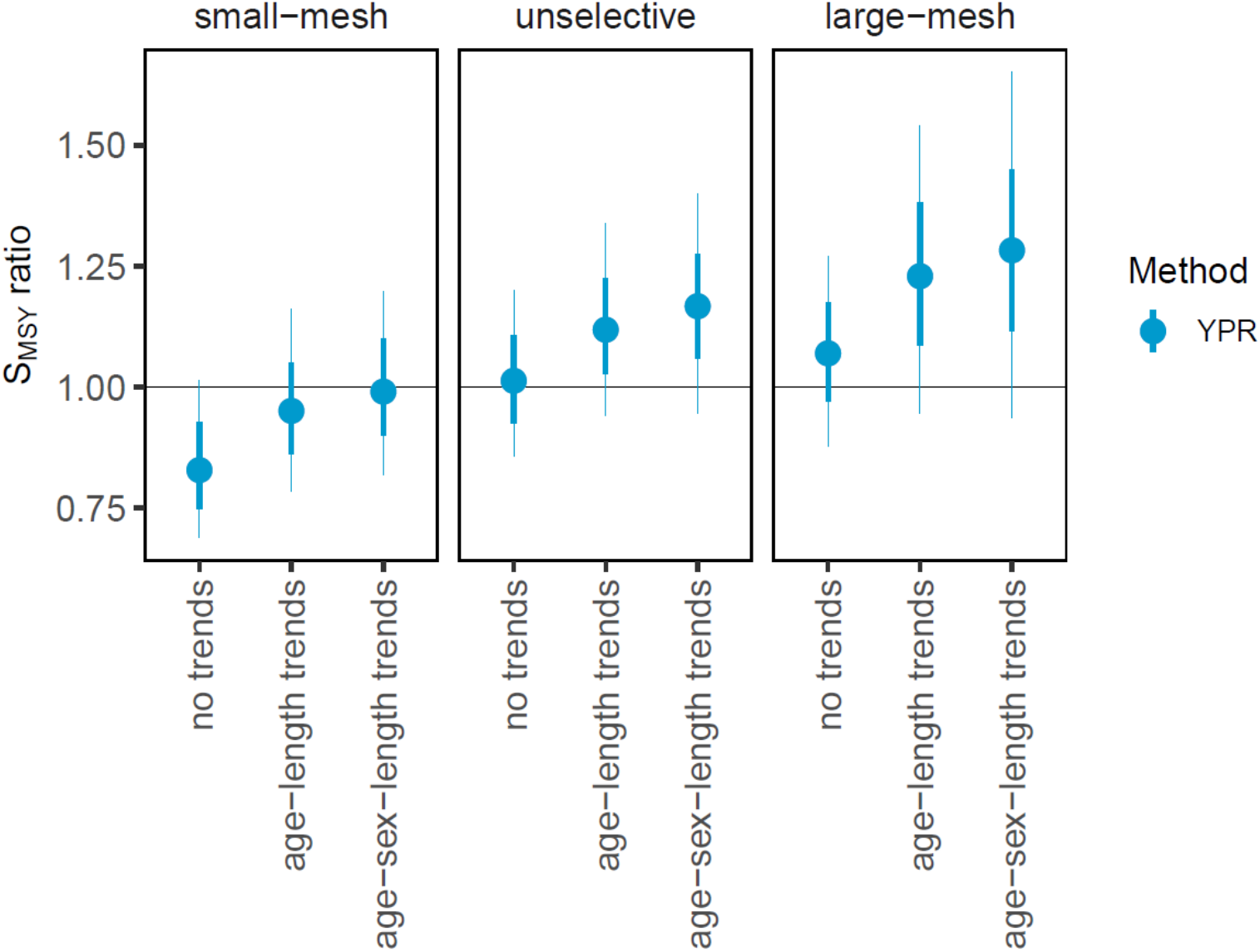
Ratio of S_MSY_ estimates for the yield-per-recruit relative to the traditional Ricker model. Shown are median values (circles) with 50% quantiles (thick bars) and 80% quantiles (thin bars) for the yield-per-recruit relative to the traditional Ricker model (YPR). S_MSY_ estimates were evaluated at the end of the historical period after 50 years of demographic trends. The horizontal line at 1.0 indicates equal estimates among the two models.

**Figure S4:**
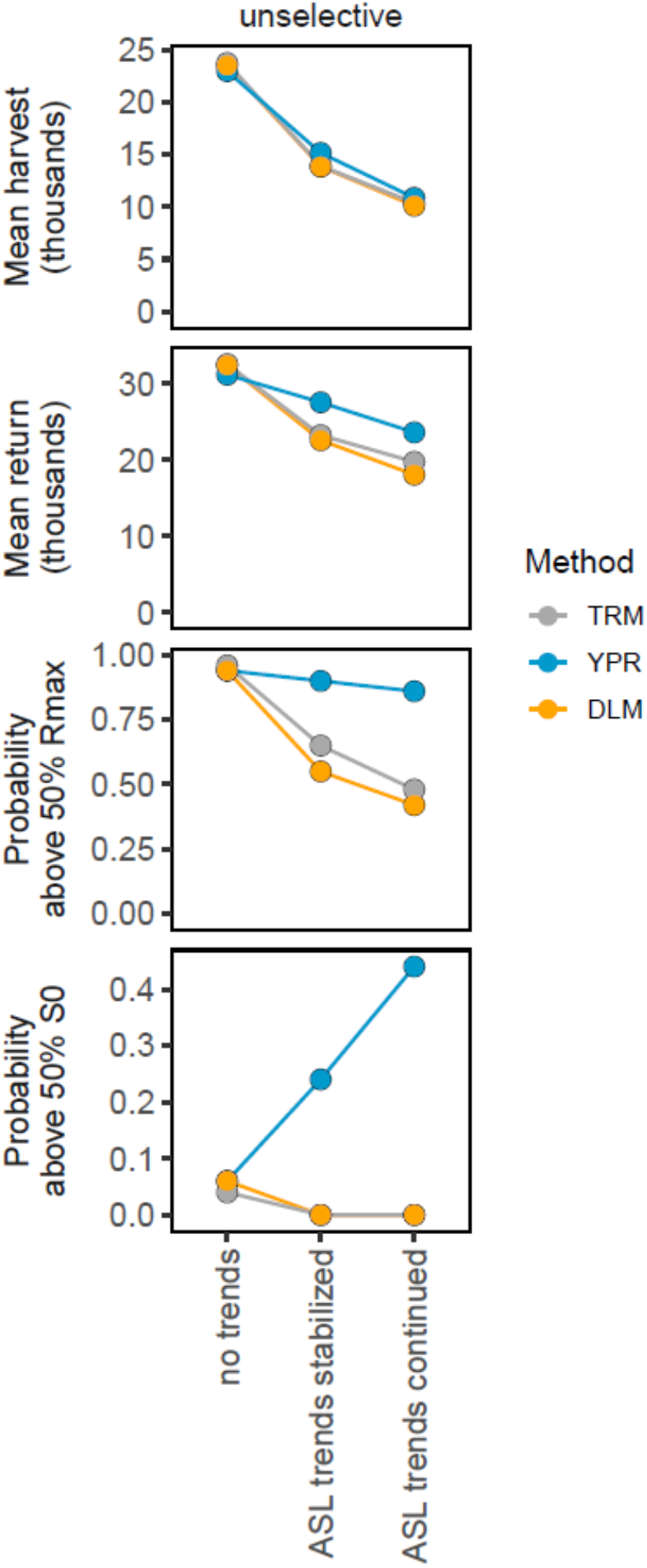
Evaluation of fishery management performance using an implementation error *σ*_*i*_ of 0.15. Shown are median values of the four examined performance metrics (rows) for each estimation method (colors) as a function of demographic trends assuming an unselective fishery. Symbology as in Figure 3 of the main text.

